# Protein diffusion on membrane domes, tubes and pearling structures

**DOI:** 10.1101/2020.10.08.331629

**Authors:** R. Rojas Molina, S. Liese, A. Carlson

## Abstract

Diffusion is a fundamental mechanism for protein distribution in cell membranes. These membranes often exhibit complex shapes, which range from shallow domes to elongated tubular or pearl-like structures. Shape complexity of the membrane influences the diffusive spreading of proteins and molecules. Despite the importance membrane geometry plays in these diffusive processes, it is challenging to establish the dependence between diffusion and membrane morphology. We solve the diffusion equation numerically on various curved shapes representative for experimentally observed membrane shapes. Our results show that membrane necks become diffusion barriers. We determine the diffusive half time, *i.e.*, the time that is required to reduce the amount of proteins in the budded region by one half and find a quadratic relation between the diffusive half time and the averaged mean curvature of the membrane shape. Our findings thus help to estimate the characteristic diffusive time scale based on the simple measure for membrane morphology.

**Significance statement:** Diffusion is an integral process for distributing proteins throughout biological membranes. These membranes can have complex shapes and structures, often featuring elongated shapes such as tubes and like a necklace of pearls. The diffusion process on these shapes is significantly different from the well studied planar substrate. We use numerical simulations to understand how the characteristic diffusion time is a function of membrane shape, where we find the diffusion of proteins on strongly curved shapes is significantly slower than on planar membranes. Our results provide a simple relationship to estimate the characteristic diffusion time of proteins on membranes based on its mean and Gaussian curvature.

## 1 Introduction

Diffusion is a fundamental transport mechanism that plays a key role in many biological processes (1; 2). Diffusion is the main mechanism for transport and mixing of components in primitive cells (3), it facilitates the formation of protein oligomers and lipid-protein assemblies associated with the metabolism and signaling in cells (4; 5; 6). It also allows the formation of protein gradients needed to establish cell polarity and trigger morphogenesis (7).

The transport of proteins and other molecules often occurs in membranes with complex shapes, which can be the result of biological process that require membrane deformation, such as endo/exocytosis (8) and communication pathways for material transport between the Golgi apparatus and the endoplasmic reticulum, requiring the formation of tube networks (9). Many effects can slow down the diffusion on biological membranes, *e.g*, cytoskeletal barriers and molecular crowding, among others (10; 11). In addition, it has been shown that the shape of the membrane itself affects how proteins or molecules move along it. Experiments reveal that the radius and length of tubes influences the time required to distribute proteins (12). The same observation was made by a theoretical approach where the diffusion equation is solved on tubular shapes, which shows agreement with the equilibration time found in experiments (13). However, biological membranes may exhibit a variety of shapes that differ significantly from a cylindrical tube: *e.g*, membranes form pits in early stages of endocytosis, and nearly spherical vesicles joined with the surrounding membrane by a narrow neck at later stages (14; 15). In addition, membranes can also form concatenated buds joined by narrow bridges or necks (16; 17; 18). Numerical simulations of diffusion on pearled structures have shown that the pearl geometry, together with diffusion barriers created by certain proteins are responsible for the sequestration of cargo in the buds (16). These results suggest that the membrane geometry has an important effect on the lateral diffusion of molecules and proteins (19; 20; 21).

The diffusion process on a curved two-dimensional surface depends on the local curvature and is thus more complex than diffusion on a flat surface. For small membrane deformations, theoretical studies have shown that an effective diffusion coefficient can be derived in terms of the surface curvature (22). A generalized theoretical treatment of the diffusion on an arbitrary surface gives a solution of the diffusion equation as an expansion around the solution on a flat surface. The coefficients are highly complex functions of the curvature, only simplified in cases where the surfaces have constant Gaussian curvature (23), which in general is not the case for biological systems. In addition, a curvature-dependent diffusion coefficient can be obtained assuming that the membrane curvature induces changes of the membrane thickness (24), introducing corrections to the diffusion coefficient of membrane inclusions predicted by the Saffman-Delbruck theory (25), which relates the diffusion coefficient with the viscosity of the surrounding media and the membrane thickness. However, these results can not be generalized to strongly curved membrane shapes such as tubes or pearl-structures.

The wide range of shapes of biological membranes makes it challenging to obtain a generalized analytical solution for the diffusion equation on a generic surface. To understand how proteins diffuse on complex membrane shapes such as domes, tubes and pearling structures we develop a mathematical model for their diffusive dynamics on these static shapes. The model is solved numerically, where the numerical simulations allow us to describe the characteristic time it takes to diffuse away from a curved region and discuss the implications of the Gaussian and the mean curvature in terms of the half-time i.e., the time for the protein density to reduce by one half in the budded region.

## 2 Methods

### 2.1 Parameterization of the membrane

The membrane shapes that we consider here are shown in Fig. 1: an Ω shape (Fig. 1a), a dome shape (Fig. 1b), pearled structures with different number *n* ∈ [2, 3, 4] of pearls (Fig. 1c) and cylindrical tubes caped by a half sphere (Fig. 1d). Each shape contains a budded region with non-zero mean curvature at the center. We denote the area of the budded region as *A*_*bud*_. The budded region continuously transitions into the outer region with zero mean curvature. A detailed description of the different shapes are presented in the Supplementary Material (SM). All shapes are parametrized by the arc-length *S* measured along the curved surface and the tangent angle *ϕ* as shown in Fig. 1a.

**Figure 1:**
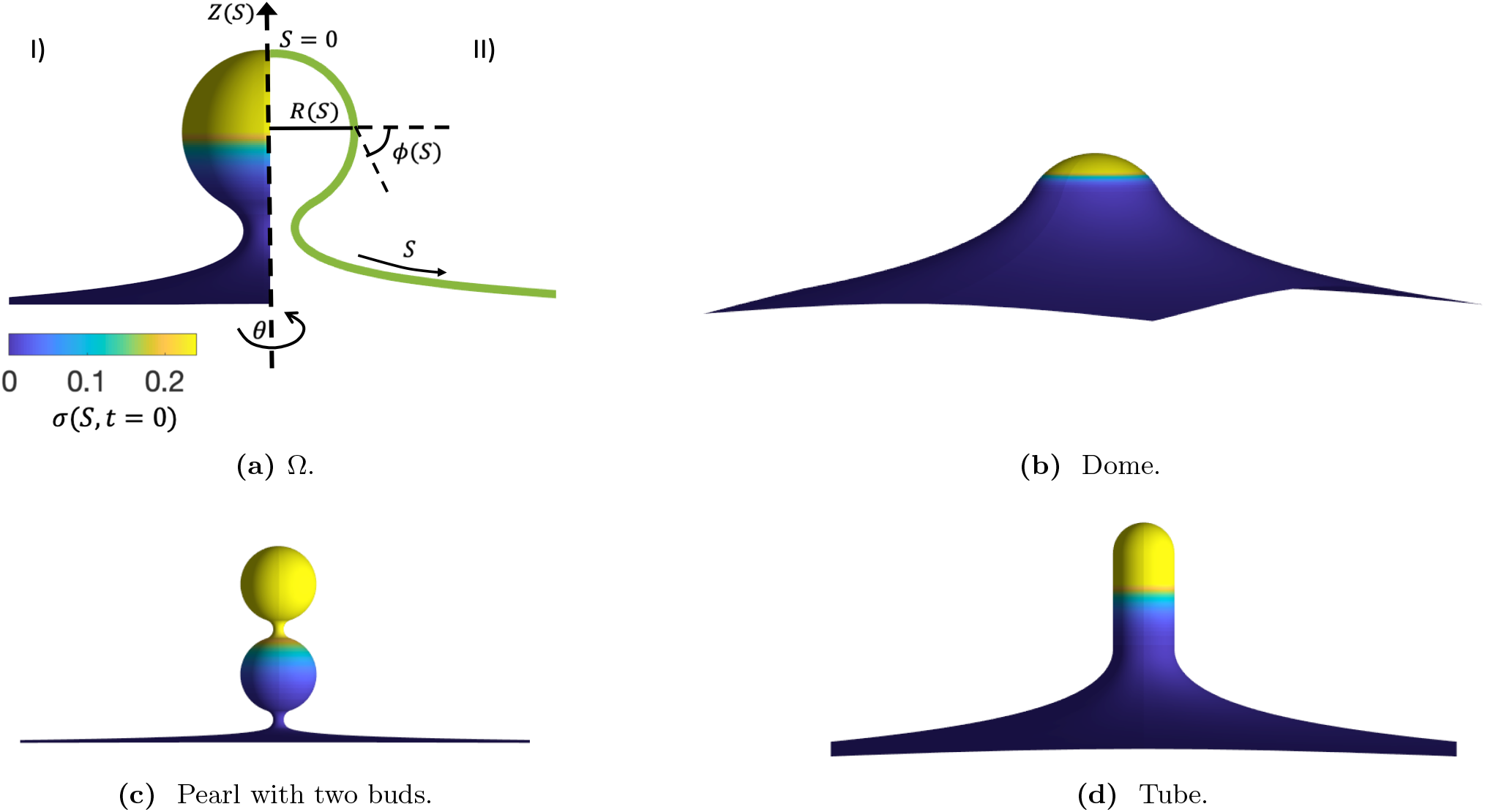
(a) I) A membrane with an Ω shape. The colorbar represents the scaled protein density *σ*(*s, t* = 0). II) The membrane surface parametrization in axisymmetric coordinates, where *S* is the arc-length measured along the membrane, *R*(*S*) is the radial coordinate, *ϕ*(*S*) is the angle that the curved membrane forms with respect to the horizontal *R*-axis and *Z* is the height of the membrane. The angle *θ* is the rotation around the symmetry axis. (b)-(d) Different membrane shapes considered here: (b) a dome shape, (c) a pearl structure with two buds and (d) a tube. Initially the total protein density *m*_*tot*_ is the same for all shapes. The shapes are axially symmetric and represent shapes commonly observed in biological membranes.

In this geometry, the variables *R*, *Z* and the membrane area *A* satisfy the following differential equations:

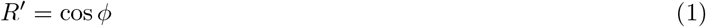

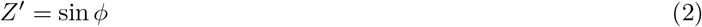

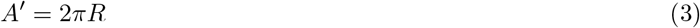

where 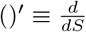. The mean curvature *H* of the membrane in the arc-length parametrization is given by (26):

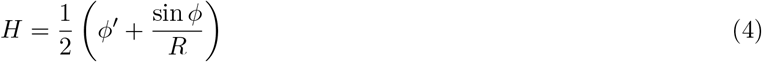

To generate the different shapes, a piecewise constant curvature radius is defined a long the membrane surface.

### 2.2 The diffusion equation

We assume now that the density of a generic protein or molecule is described by a continuous field 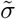 that depends on the arc-length *S* and the time *t*, 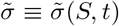. The field 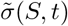 follows the diffusive dynamics on the different shapes shown in Fig. 1. The diffusion equation in the absence of sources, under axial symmetry is given by (26):

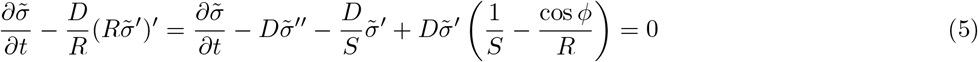

where *D* is the diffusion coefficient and assumed to be constant. On a flat membrane, with *ϕ* = 0 and *R* = *S*, Eq. 5 is written as 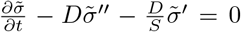. On a curved surface the diffusion equation (Eq. 5) contains an additional term 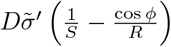, which depends both on the local curvature and the gradient of the protein density. Diffusion on a curved surface is as such different from the diffusion on a flat surface and the combined effect of curvature and protein distribution can in general not be captured in a single effective diffusion constant.

We write the diffusion equation (Eq. 5) in dimensionless form using the square root of the area of the budded region of the membrane, 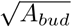 (see the SM) as a characteristic length. Then the time scale is given by *τ* = *A*_*bud*_*/D* and the dimensionless variables are 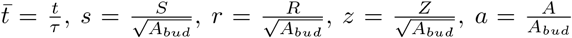 and 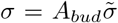. By introducing these scaling relations into Eq. 5 we obtain:

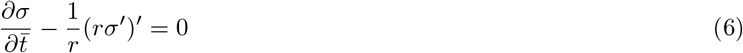

As an initial protein density, we consider *σ*(*s, t* = 0) = *c*(1 −tanh [5(*a*(*s*) − *a*_0_)]) for all membrane shapes, where there is a one-to-one correspondence between the arc-length *s* and the area *a*(*s*), given by Eq. 3. The constants *c* and *a*_0_ are chosen such that *σ*(*a*(*s* = 0)) = 0.5 and *m*_*tot*_ = 0.24, with 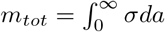 the total amount of proteins. As shown in Fig. 1, we consider an initial protein distribution that is nearly homogeneous in the center of the membrane and that vanishes at the outer, flat membrane region. As time proceeds, this initial density will diffuse over the entire membrane surface. The boundary condition imposed on the diffusion equation is such that there is no flux of protein at the boundaries of the spatial domain, *i.e*, *σ*′(*s* = 0) = 0 and *σ*′(*s* → ∞) = 0. In this case, the total quantity of protein *m*_*tot*_ is conserved over time, as the proteins cannot escape from or flow through the membrane boundaries. Eq. 6 is integrated numerically in time with a time step *dt* = 5 × 10^*−*3^ to obtain the spatio-temporal density profile on the membrane, using a implicit time discretization and a centered difference scheme to solve the spatial derivative of the protein density.

## 3 Results

In Fig. 2, the protein density *σ* is shown as a function of the membrane area *a* at different points in time, 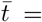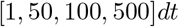. These density profiles exhibit notable qualitative differences. The Ω shape and the pearl-like shapes exhibit a staircase-like protein profile in the budded region (*a* < 1), where the protein density approaches a nearly homogeneous distribution within the individual pearls. In contrast, neither the dome nor the tubular shape show a step-like protein distribution at any point in time. Comparing, dome/tubular shapes and Ω/pearl-like shapes at the latest time point, we find for the latter that the protein density in the budded region is much higher, indicating that the presence of narrow necks slows the diffusion of the proteins.

**Figure 2:**
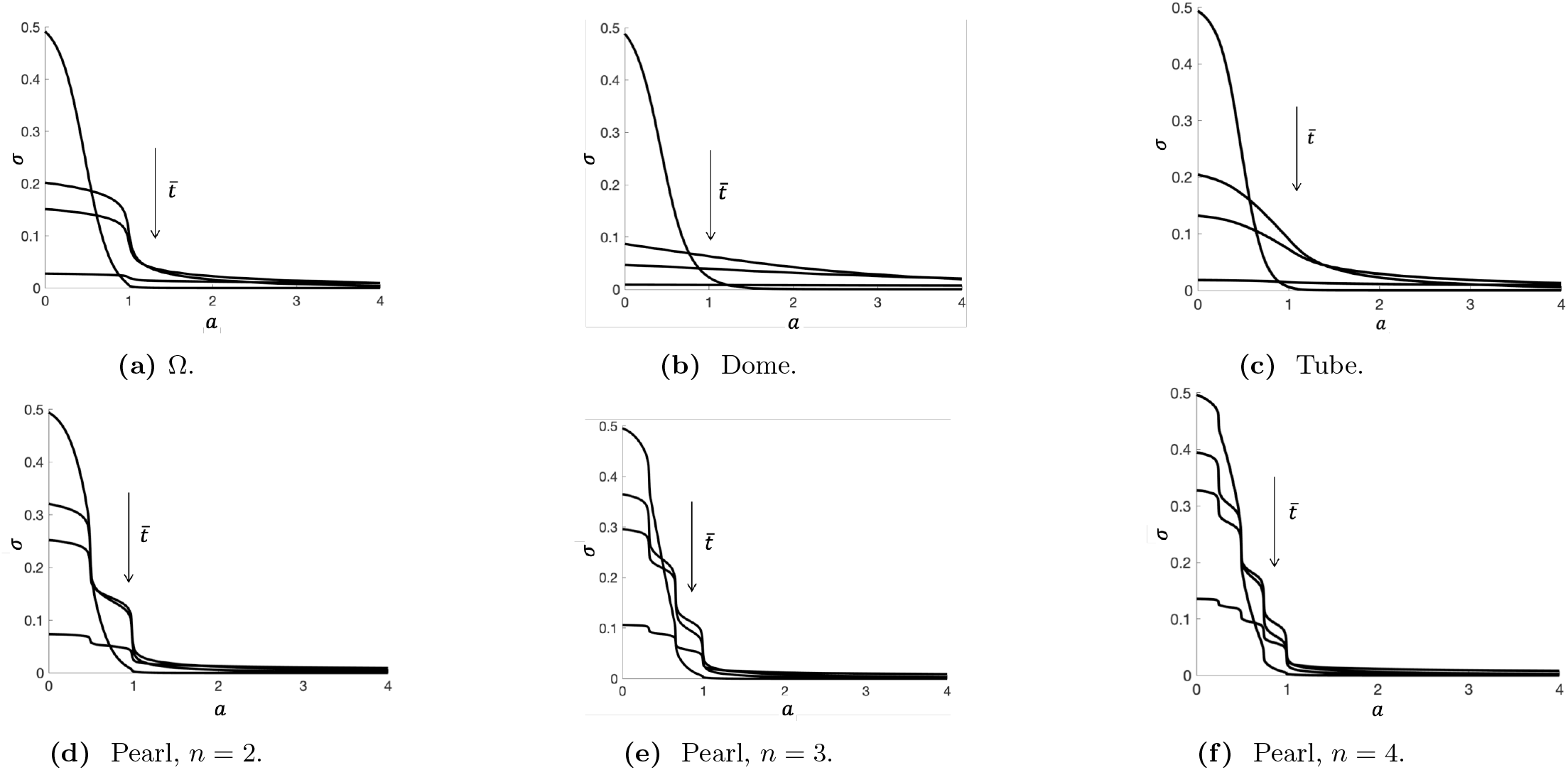
Density profiles as a function of the membrane area *a* at different times, 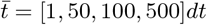, for (a) an Ω shape, (b) a dome shape, (c) a tube, and (d)-(f) pearling structures with different number of pearls. As time proceeds, the initial protein density spreads over the membrane area, where the profiles are qualitatively different. In the case of the Ω (a) and the pearling shapes ((d)-(f)) the protein density has a staircase-like profile, along each of the pearls, indicating that the narrow necks prevents the profile of having a smoother transition. The overall effect of the membrane necks on the diffusion on buds and pearls is to slow down the exit of proteins from the budded structure. In all simulation, the initial density is given by 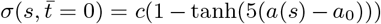, where *c* and *a*_0_ are such that 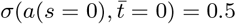 and *m*_*tot*_ = 0.24.

To characterize the effect of the membrane shape on the diffusive dynamics by a single parameter, we define the amount of proteins in the budded region 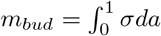 and determine how *m*_*bud*_ evolves in time. In Fig. 3a we show the time evolution of *m*_*bud*_ on the pearling structure with *n* = 2, the Ω shape, the tube and the dome shape. Our simulations show that, in accordance with the qualitative discussion of the density profiles in Fig. 2, the amount of proteins in the budded region decreases faster in shapes that do not have narrow necks, such as the dome shape and the tube, while *m*_*bud*_ decreases slower on the Ω shape and the pearling structure. In addition, Fig. 3b shows *m*_*bud*_(*t*) for the different pearl-like structures, with *n* = 2, 3, 4, where we see that *m*_*bud*_ decreases slower with larger number of pearls.

**Figure 3:**
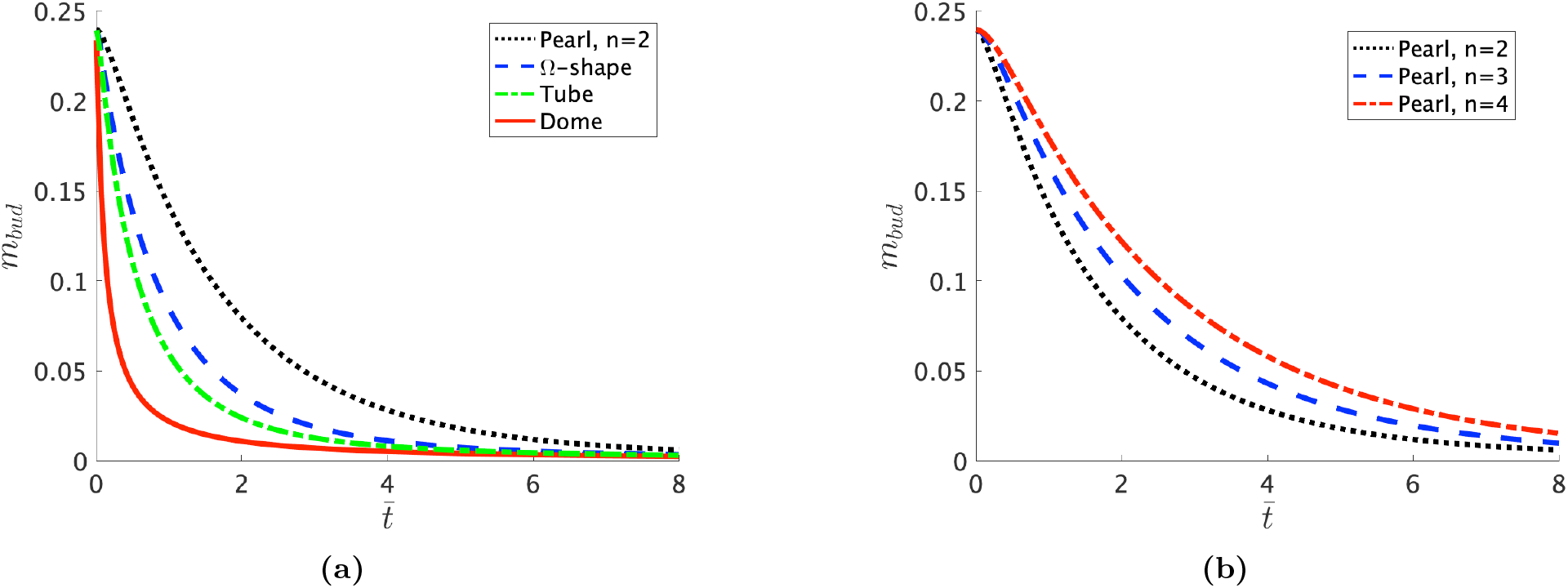
The total protein density on the budded region 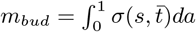 as a function of time for different membrane geometries. (a) 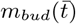 on the dome, tube, Ω and pearl-like (*n* = 2) shape. (b) The effect of the number of pearls on the pearl-like structure on 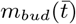. The proteins exit the pearled region slower if the number of pearls, *n*, increases.

The results in Fig. 3 highlight that the membrane shape influences both the temporal and spatial evolution of the protein distribution. To characterise the diffusion process by a single quantity, we define 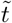 as the half-time, *i.e*, the time it takes for the total amount of proteins in the budded region to be reduced by 50%. On a flat membrane the half-time is denoted as 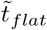. The ratio 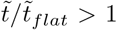 on all budded membranes, indicating that diffusion on a flat surface is faster compared to diffusion on a curved surface.

We aim at relating 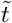 to the characteristics of the membrane shape where we consider the Gaussian curvature *K* averaged over the budded area 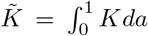, the Gaussian curvature in the membrane neck, *K*_min_, *i.e.*, where the membrane radius is minimal and the mean curvature averaged over the budded area, 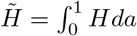.

In Fig. 4a we plot 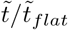 as a function of 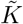, and observe that 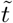 is not clearly correlated to the averaged Gaussian curvature 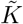. 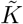 is similar for the tubes and the single bud, but their 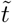 is different. All the pearled structures, on the other hand, have a similar 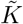, but 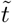 is also clearly different different and up to five times larger with respect to 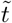 in the tubes and in the single vesicle. 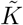 for the dome is clearly different from the 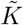 of the tubes and the pearls. However, the diffusion time of buds is similar to tubes. Fig. 4b shows 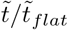 as a function of the minimum value of the Gaussian curvature, *K*_min_, which corresponds to the point along the arc-length *s* where the neck is smallest. The pearling shapes with a large number of vesicles have a smaller *K*_min_. In contrast, the shapes where the neck regions are absent, as in the domes and the tubes, have a smaller and close to zero *K*_min_. Fig. 4b indicates that 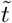 increases as the membrane have more constricted regions, which creates a clogging effect.

**Figure 4:**
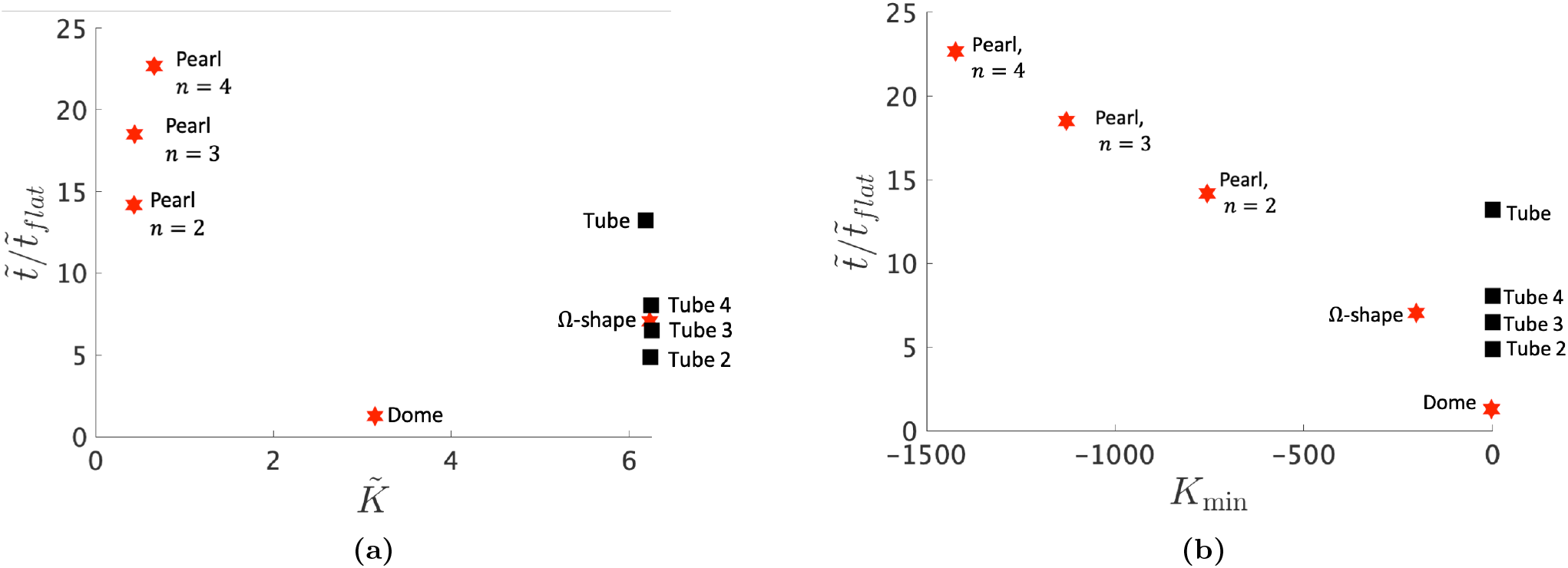
(a) The ratio 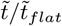 as a function of the averaged Gaussian curvature 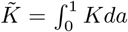. There is not a clear dependence between 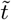 and 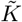 across all the shapes considered. The tubes and the Ω-shape have a similar 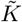, which is larger than the 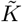 of the pearled structures. The time 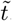, however, is clearly different, specially in the pearled structures, which also have a similar 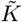. (b) The ratio 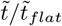 as a function of the minimum value of the Gaussian curvature along the arc-length, *K*_min_, for all the shapes considered. *K*_*min*_ corresponds to the point where the neck joining the pearl with the surrounding flat membrane is minimal, in the case of the pearls and the Ω-shape. In the tubes, *K*_*min*_ corresponds to the tube rim, and in the dome *K*_*min*_is the point beyond which the mean curvature vanishes.

In Fig. 5 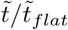 is shown as a function of the averaged mean curvature 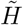 in logarithmic axis. We find an approximately quadratic relation between 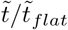 and 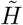, where the diffusion on tubular shapes is slightly faster, *i.e.* lower 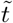, compared to pearling structures. An analytic expression for 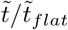 can in general not be derived. However, we can compare the effective diffusion constant on simpler shapes, namely on an infinite cylinder and a sphere with our numerical results, which capture the main characteristics of the diffusion process. Holyst et al. found the following expression for the effective diffusion constant *D*_eff_ of a single particles diffusing on an infinite cylinder and a sphere, respectively, 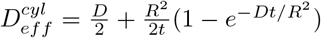 and 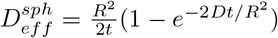 (27), with *R* the radius of the cylinder or the sphere. The effective diffusion constant is defined as *D*_*eff*_ =< *x*^2^(*t*) > /4*t*, with < *x*^2^(*t*) *>* the mean squared displacement of the particle. In both cases the diffusion constant is time-dependent and decreases over time. Furthermore, the diffusion process is slowed down, *i.e. D*_*eff*_ < *D*, for both the cylinder and the sphere, and the effective diffusion constant on a sphere is always smaller then the effective diffusion constant on a cylinder, given that both shapes exhibit the same radius.

**Figure 5:**
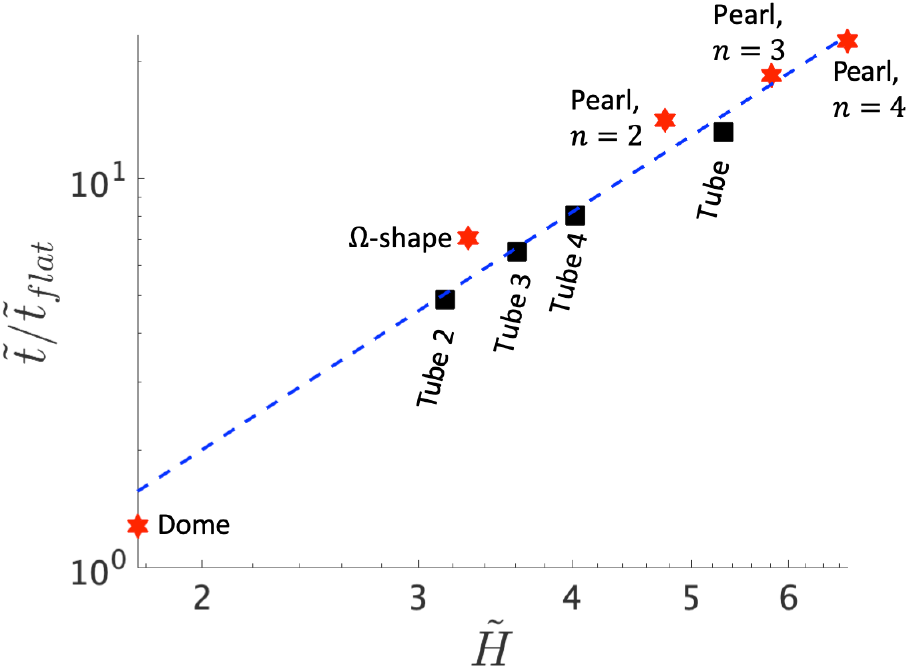
The ratio 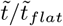 as a function of the averaged mean curvature 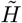 for each of the shapes considered, in logarithmic scale. The blue dashed line represents a fit 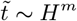, where *m* = 2.03 is the average of the slopes of the logarithmic relation between 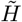 and 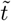, indicating that the exit time follows aroximately a quadratic relation with respect to 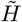. However, some of the pearled structures do not follow the quadratic fit. This difference can be rationalized qualitatively associating the tubes with a cylinder of infinite height and the shapes with spherical buds with a sphere. In these simpler geometries, the diffusion equation can be solved analytically, and an effective diffusion coefficient *D*_*eff*_ can be defined. On a cylinder with the same radius of a sphere, *D*_*eff*_ is larger as time proceeds, indicating than in cylindrical geometries the diffusion is faster. Here, we observe that in general, 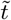 is smaller in the tubes, indicating a faster diffusion with respect to the pearls and the Ω-shape. However, there are other effects that play an important role on the diffusion from these structures, such as the obstacles produced by the neck regions in the Ω-shape and the pearled structures, and also the obstacles to diffusion in the region joining the budded structure with the surrounding, flat membrane.

These analytical results indicate that the effective diffusion on a cylinder is faster than in a sphere as time proceeds and help to understand why 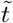 is smaller in the tubes than in the dome and the pearled structures, as shown in Fig. 5. As shown in Fig. 5, the radius of the tube (and hence the mean curvature 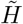) is correlated to 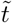, indicating that the tube radius has an effect on the exit of protein from the tubular region towards the flat surface (13) as a result of the steep change in the derivative of *r*, as it passes from being *r*′ = 0 (*r* constant) in the flat region of the tube to an increasing function of the arc-length *s* (*r*′ = 1). This sharp transition is more abrupt when the tube radius is smaller, causing a larger 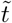when the mean curvature of the tube increases. In the case of the dome and the pearls, the diffusion is not limited to a single sphere, as the proteins are free to diffuse outside each bud, though the process is slowed significantly.Finally, 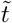 is a time scale at which all the effects mentioned above play an important role on the diffusion of proteins from the different shapes, as a significant part of the initial protein density have exit the deformed regions and gone thought the obstacles imposed by the membrane geometry.

## 4 Conclusions

In this work we study the influence of characteristic shapes found in biological membranes; Ω-shaped buds, pearls, domes and tubes, on the diffusion of a protein density field. The shapes considered have a strong influence on the density profiles obtained as time proceeds, and also influence the characteristic time after which the proteins exit the budded region: the presence of narrow necks (pearls, Ω-shaped buds) prevents a fast decay of the total density on the budded structure, as compared to the shapes where such necks are absents (tubes, domes). To characterize more precisely the effect of the shape on diffusion, we have determined the averaged Gaussian and mean curvature for each shape. The time associated with proteins leaving the budded structures, 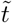, does correlate with the mean curvature, but not with the Gaussian curvature: while 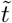 follows a quadratic relation with respect to 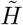, it does not show a clear dependence with respect to 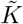. However, a larger 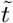 is associated with a smaller Gaussian curvature *K*_min_, in the neck region. This indicates that constricted regions indeed delay the exit of proteins from curved membranes.

The relations between 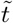, 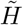 and *K*_min_ can help to estimate relevant time scales related to the diffusive motion of proteins on complex membrane shapes. Our results suggest that the exit of proteins is strongly affected in elongated shapes, such as the tubes and the pearled structures. These structures are observed in different biological contexts and in some cases can be reproduced by different experimental techniques. In addition, photobleaching provides a powerful technique to investigate the mobility of proteins inside cells (28) and in the membrane surface (29). This technique have allowed to corroborate experimentally the recovery time predicted theoretically in tubular geometries (13), and motivates the use of theoretical models to predict diffusive behavior of proteins and molecules in more generic shapes.

## 5 Sulementary Material

### 5.1 Membrane shape geometry

Each shape contains two regions, the inner budded region and the outer region with zero mean curvature. We define the second principle curvature in the inner region as follows:

**Dome and** Ω **shape**:

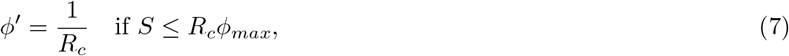

with *ϕ*_*max*_ = *π/*3 for the dome shape (Fig. 6b) and *ϕ*_*max*_ = 5*π/*6 for the Ω shape (Fig. 6a). In case of a dome shape we consider the region with non-zero mean curvature as the budded region. In case of an Ω-shape we denote the membrane portion above the minimal neck radius as the budded region. The radius *R*_*c*_ is adjusted such that the area of the budded region is equal to 1.

**Pearl (n=2)**:

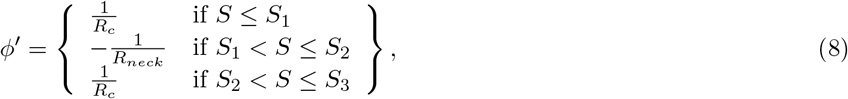

where *S*_1_ = *R*_*c*_*ϕ*_*max*_, *S*_2_ = *S*_1_ + (2*ϕ*_*max*_ − *π*)*R*_*neck*_ and *S*_3_ = *S*_2_ + (2*ϕ*_*max*_ − *π*)*R*_*c*_ and *ϕ*_*max*_ = 9*π/*10 (Fig. 7a). The ratio between *R*_*c*_ and *R*_*neck*_ is set to *R*_*neck*_ = *R*_*c*_/4. More pearls can be added to the shape by defining additional neck and pearl regions.

**Figure 6:**
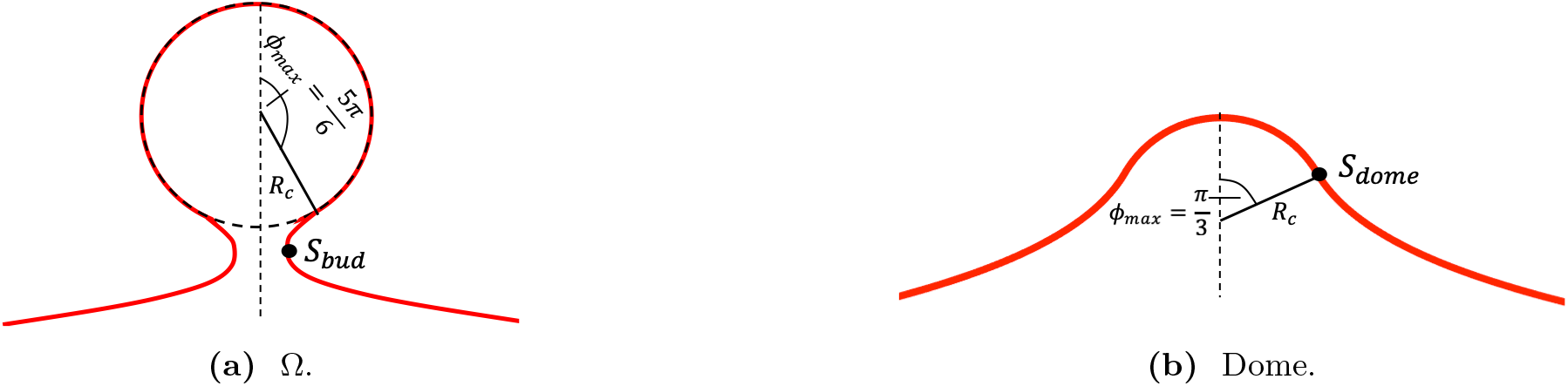
(a) A schematic representation of the Ω shape (a) and dome shape (b). The value of the coordinate *S* at which the radial distance to the symmetry axis *R* is minimal, *S*_*bud*_ defines the neck of the Ω shape. The neck of the dome shape is defined at *S*_*dome*_. In the outer region the membrane has a vanishing mean curvature, *H* = 0

**Figure 7:**
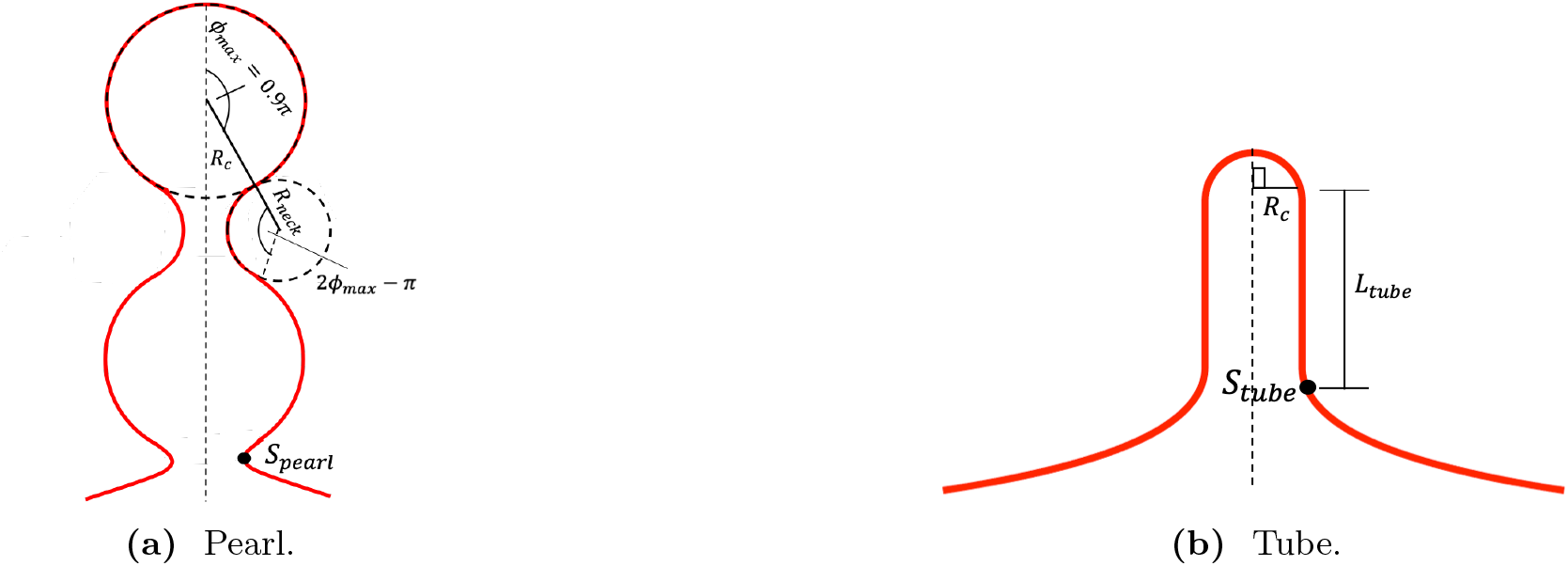
(a) A schematic representation of a pearled structure. It is assumed that the bud and the neck have radius of curvature *R* and *R*_*neck*_, respectively, over regions spanned by the angle *ϕ*_*max*_ and 2*ϕ*_*max*_ − *π*. In the region outside the pearl, the membrane has vanishing mean curvature, *H* = 0, which means it approaches a flat surface outside the budded region. The coordinate *S*_*pearl*_ defines the area of the budded structure and corresponds to the minimal radial distance *R* joining the pearl with the rest of the membrane. (b) A schematic representation of a tubular structure. The hemisphere region at the top of the tube has a radius of curvature *R*_*c*_, spanned by the angle 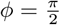. The tube has a height *L*_*tube*_ measured from the bottom of the hemisphere to the tube rim. In the region outside the tube, the membrane has vanishing mean curvature, *H* = 0.

**Tube**:

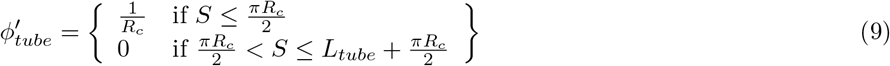

The length *L*_*tube*_ are adjusted such that the adimensional area of the budded region is equal to 1. In the outer region, the second principle curvature is given by:

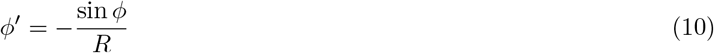

for all shapes to achieve a vanishing mean curvature

### 5.2 Conservation of the total protein density

We consider a diffusive process where the boundary conditions are such that there is no flux of proteins at the boundaries of the spatial domain. This means that there are not proteins entering or leaving the membrane. Under these conditions, the total amount of proteins *m*_*tot*_ = ∫ *σda*, must be constant in time. In our simulations, we have set as *m*_*tot*_ = 0.24 at 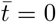. In Fig. 8 we show *m*_*tot*_ as function of time, where, as predicted, *m*_*tot*_ is constant over time on the bud. Exactly the same plot is obtained on all the other shapes considered, and for the sake of brevity we do not show all of them.

**Figure 8:**
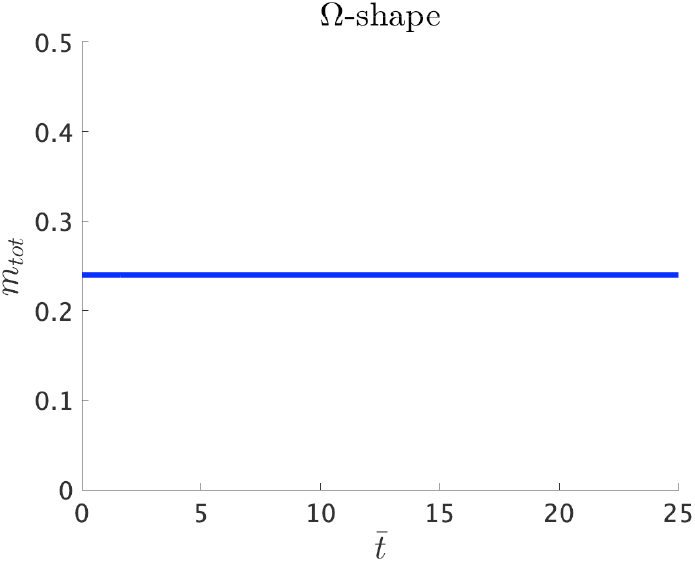
The total amount of proteins *m*_*tot*_ on the bud as a function of time. Consistent with the zero flux boundary condition imposed to solve the diffusion equation on the different shapes, *m*_*tot*_is conserved over time.

### 5.3 Sensitivity to different initial density profiles

As mentioned in the main section, we have chosen a initial density profile of the form 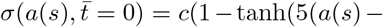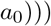, where *c* is adjusted in such a way that the value of 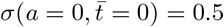 and *a*_0_ is adjusted to give a total amount of proteins *m*_*tot*_ = 0.24. However, different values of 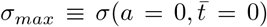 yield different values of *a*_0_, so we can consider various initial density profiles that gives the same *m*_*tot*_. In Fig. 9 we show different initial density profiles, 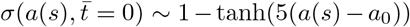 and the density profiles obtained with each of these initial conditions at later time steps on the Ω shape (Figs. 9b and 9c), the dome (Figs. 9e and 9f), the pearl structure with *n* = 3 buds (Figs. 9h and 9i) and the tube (Figs. 9k and 9l) showing that as time proceeds the effect of having different initial condition vanishes, as the density profiles tend to become equal regardless of the chosen initial profile. In the case of the pearl structure, a longer time is needed for the profiles to become equal.

**Figure 9:**
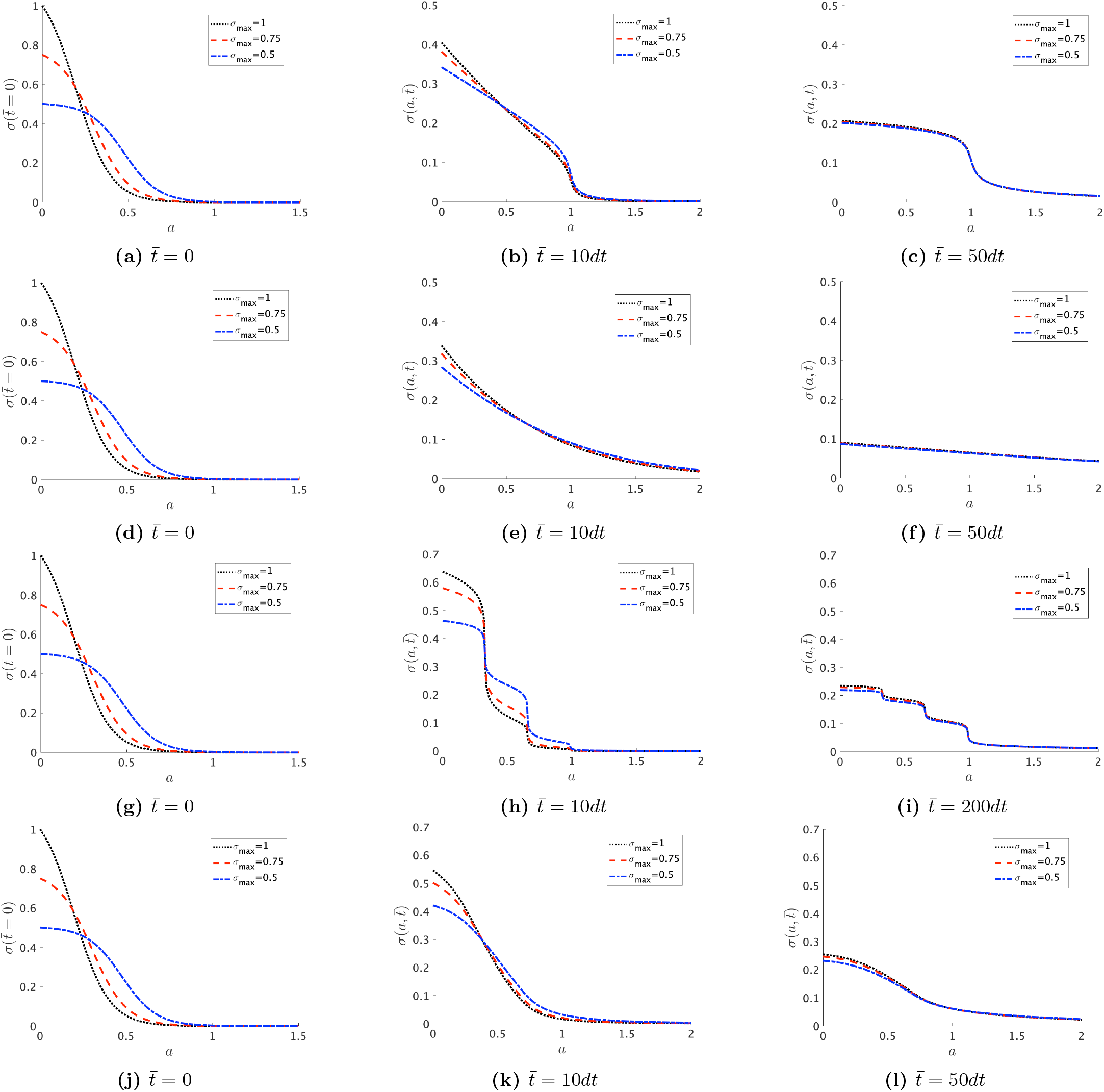
Three different initial protein profiles, 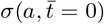 profiles with σ_max_ = 0:5; 0:75; 1:0, that yield the same total amount of proteins *m*_*tot*_ = 0:24, for different shapes: ((*a*) - (*c*)) Ω-shape, ((d)-(f)) dome, ((g)-(i)) pearl with n = 3 buds and ((j)-(l)) a tube. At later times 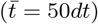 the density profiles in the Ω-shape, the dome and the tube tend to become equal, regardless of the chosen initial condition, but in the case of the pearl structure, a longer time 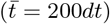 is needed for the profiles to become equal.

Different initial conditions lead to different density profiles during the first time steps of the simulations, as shown in Fig. 9, but the initial conditions have no effect on the amount of proteins in the budded region *m*_*bud*_ as time evolves, as shown in Fig. 10, indicating that the diffusion of proteins away from the budded region does not depend on the initial condition.

**Figure 10:**
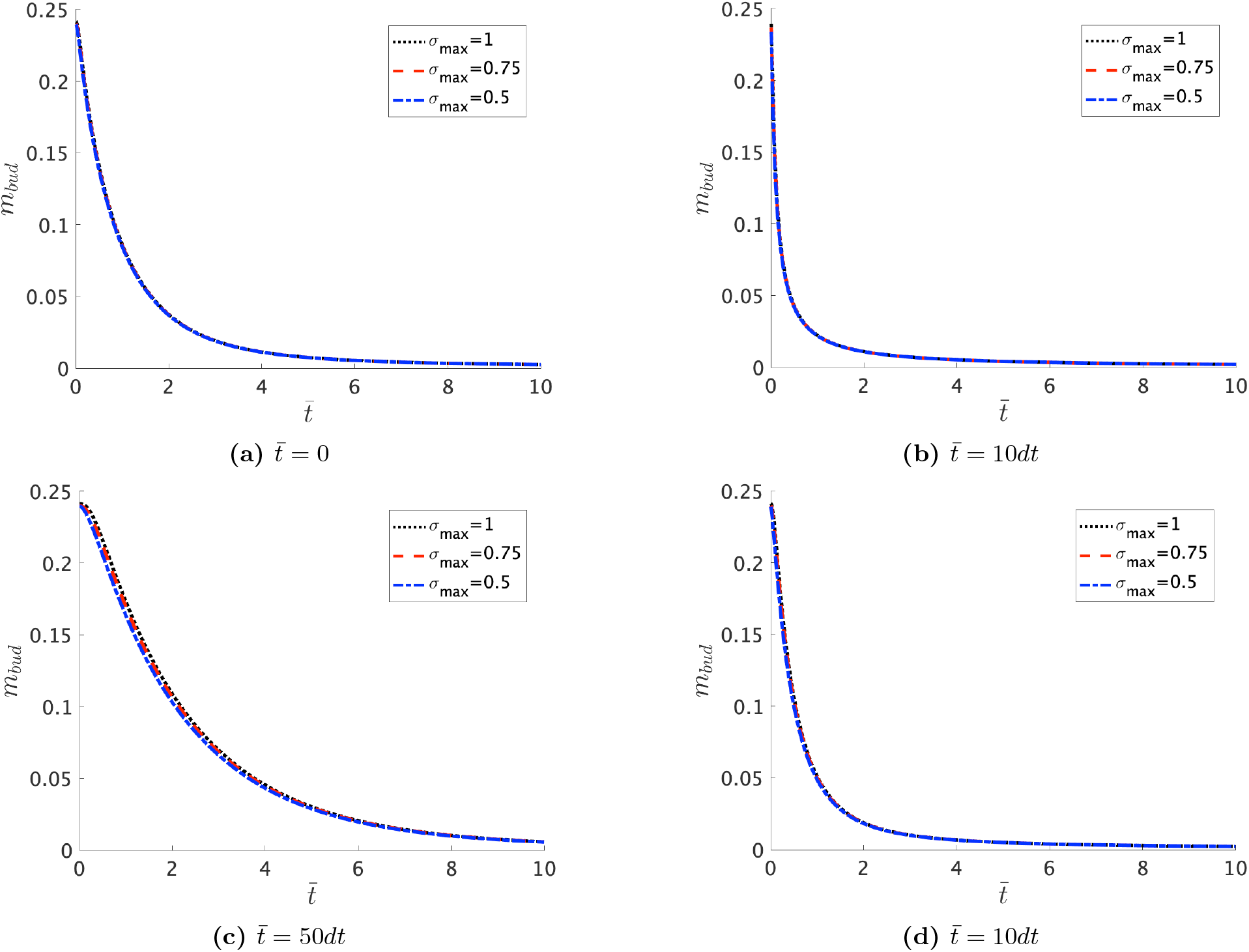
The total amount of proteins in the budded region *m*_*bud*_ as a function of time, for the initial conditions shown in Fig. 9a and for different shapes: (a) Ω-shape, (b) dome, (c) pearl with *n* = 3 and (d) a tube. These initial conditions have no effect on the time evolution of the protein density on the budded region, as the different curves are overlaing.

